# Flavor Classification/Categorization and Differential Toxicity of Oral Nicotine Pouches (ONPs) in Lung Epithelial Cells

**DOI:** 10.1101/2022.07.06.498919

**Authors:** Sadiya Shaikh, Wai Cheung Tung, Joseph Lucas, Shaiesh Yogeswaran, Dongmei Li, Irfan Rahman

**Affiliations:** Department of Environmental Medicine, University of Rochester Medical Center, Rochester, NY, USA; Department of Clinical & Translational Research, University of Rochester Medical Center, Rochester, NY, USA

**Keywords:** Cytotoxicity, Inflammation, Oxidants, Cytokines, lung epithelial cells, Oral Nicotine Pouches, Smokeless Tobacco

## Abstract

The prevalence of flavored tobacco product usage amongst youth in the United States is partly due to the emergence of non-combustible nicotine-containing products (NCNPs), including oral nicotine pouches (ONPs) and smokeless tobacco products. ONPs are available in various different flavors (mint, fruity, tobacco, dessert, citrus, coffee, wintergreen, and berry) and may use either Tobacco-Derived Nicotine (TDN) or Tobacco-Free Nicotine (TFN). Currently, several brands of ONPs are sold in the U.S and comprise a significant portion of NCNP sales in the U.S. There is a growing concern that flavored ONPs may not only induce oral health effects, but may also induce systemic toxic effects due to nicotine and other ONP byproducts being absorbed into systemic circulation through the oral mucosa. These byproducts can act locally on other tissues and may potentially cause redox dysregulation and heightened inflammatory responses systemically in the respiratory, cardiovascular, and/or renal systems. Hence, we determined the effects of flavored ONPs from four of the most widely sold brands in the U.S in inducing toxicological effects on the respiratory epithelium. Prior to analyzing the effects ONPs, we first classified ONPs sold in the US based on their flavor and the flavor category to which they belong to using a wheel diagram. Subsequently, using human bronchial epithelial cells (16-HBE and BEAS-2B) exposed to extracts of flavored ONPs, we assessed the levels of ONP-induced inflammatory cytokine release (IL-6 and IL-8), cellular Reactive Oxygen Species (ROS) production, and cytotoxicity in the airway epithelium. Our data showed that cells exposed to the lowest concentration treatments showed increased cytotoxicity, differential cellular ROS production, and proinflammatory cytokine release. The most striking response was observed among cells treated with the spearmint ONP, whereas ONPs containing original tobacco and fruity flavors showed varied levels of ROS release in 16-HBE cells. Our data suggest that flavored ONPs are unsafe and likely to cause systemic and local toxicological responses during chronic usage. Our study is a part of ongoing efforts to use *in vitro, ex-vivo, and in vivo* systems to understand how the usage of various flavored ONPs may cause both oral and pulmonary toxicity, and impact human periodontal health.

## INTRODUCTION

According to the 2021 National Youth Tobacco Survey (NYTS), approximately 2.55 million middle and high-school students in the United States (U.S.) reported current usage of tobacco products: specifically, 2.06 million (13.4%) high school students and 470,000 (4.0%) middle school students [1]. The prevalence of tobacco-product usage amongst youth in the U.S. is due to the emergence of non-combustible nicotine-containing products (NCNPs), like Electronic Nicotine Delivery Systems (ENDS) or e-cigarettes, smokeless tobacco, and nicotine pouches in the last decade [1]. Most alternative tobacco products sold within the U.S. contain nicotine; in addition to being highly addictive, nicotine is known to cause injurious responses in the lungs, heart, and kidneys [2]. Regarding youth, studies have shown that nicotine exposure in adolescence induces effects lasting until adulthood; this includes emotional dysregulation and decreased cognitive functioning [3]. An NCNP of growing concern in the U.S. is the Oral Nicotine Pouch (ONP) [1]. Among students surveyed in the 2021 NYTS, 17.2% had frequently used ONPs [1]. Like Snus (an smokeless tobacco product), ONPs are pouch-based nicotine products, products relying on the absorption of nicotine into the oral mucosa [4]. Unlike Snus, ONPs contain no components of the tobacco plant’s leaves, stem, or dust. To further explain, while some ONPs may contain tobacco-derived nicotine (TDN), they lack any other components of the tobacco plant [5]. However, like Snus, ONPs can come in various flavors (mint, fruity, tobacco, citrus, coffee, wintergreen, and berry), as represented in **Figure 1**. Moreover, the availability of this multitude of flavors amongst ONPs contributes to the prevalence of ONP usage in the U.S. [6]. We have recently identified, via Reddit social media posts, the prevalence of positive attitudes towards ONPs among Reddit/topic-discussion threads focusing on the usage of NCNPs [7]. Social media platforms like Reddit serve as platforms where users of ONPs and other NCNPs can actively share and discuss their experiences with different products, which is a significant factor in influencing attitudes and consumer behaviors revolving around ONPs [7].

**Figure 1:**
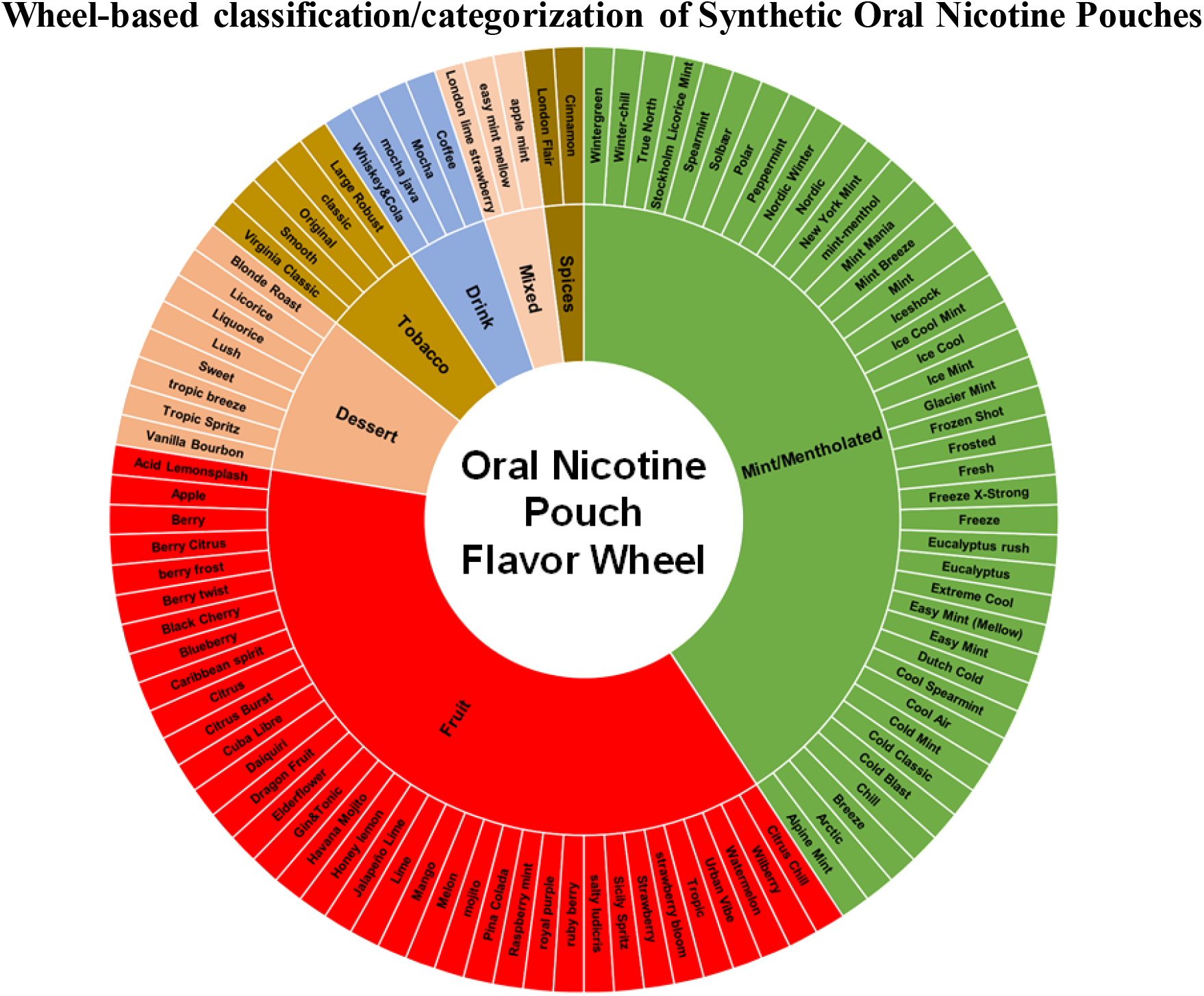
Wheel-based classification/categorization of Synthetic Oral Nicotine Pouches commonly sold in the US. The nicotine concentration of all smoke-free nicotine-based pouches ranges from 3 mg to 8 mg per pouch; mint/menthol and fruit are two of the most widely sold flavors in the US. The flavors of each pouch product in the diagram are color-coded by flavor category. The inner wheel represents the most common flavors, and the outer wheel represents specific flavors.

Regarding federal regulations on ONPs, on March 15, 2016, the FDA finalized the “Deeming Tobacco Products To Be Subject to the Federal Food, Drug, and Cosmetic Act” (the “Deeming Rule”) [8]. Under the deeming rule, the FDA’s regulatory authority over tobacco products was extended to all products containing TDN, including ONPs [8]. Subsequently, manufacturers of ONPs containing TDN were subject to pre-market assessment, required to submit specific product information to the FDA, and required to comply with marketing restrictions [9]. ONPs may contain synthetic nicotine instead TDN. However, as of April 14, 2022, the FDA’s regulatory authority was extended to include ONPs utilizing synthetic nicotine or Tobacco Free Nicotine (TFN) [10]. To explain, on March 15, 2022, the definition of a “tobacco product” under the Federal Food, Drug, and Cosmetic Act was amended to “any product made or derived from tobacco or containing nicotine from any source, that is intended for human consumption” [10]. Any ONP, regardless of whether it uses TFN or TDN, which has not submitted a pre-market application (PMTA) to the FDA, will be removed from the market in the U.S. [8, 10]. In the U.S, the sales of nicotine pouch sales increased from 163,178 units ($709, 635) in 2016 to 45,965,455 units ($216,886,819) by the end of June 2020 [11]. Moreover, from 2020-to-2021, shipments of nicotine pouches to the U.S. from Zyn (manufactured in Sweden) increased by more than 50% [12].

Despite the significant increases in the usage of ONPs in the U.S., limited studies have been conducted so far that have delved into understanding the health effects of ONP usage [4, 12–14]. However, regarding usage of smoke-free nicotine pouch-based products, studies have shown that regular usage of smokeless tobacco is associated with a higher risk for Parkinson’s disease, cancer, birth defects, type 2 diabetes, oral submucosal fibrosis, and cardiovascular disease [15, 16]. Regarding studies on the health effects of Snus and ONPs, limited studies and case reports have focused on investigating the systemic-oral-pulmonary health risks of regularly using these products [17–21]. While smokeless nicotine-based products, including Snus and ONPs, are not inhaled through the lungs, the nicotine, flavoring chemicals, and byproducts within those pouches can be absorbed across the buccal membrane into the systemic circulation; these byproducts can act locally on other tissues within the body; some of these responses are related with the cardiopulmonary system via microvasculature, liver, kidneys, the pancreas, and the esophagus [18–24]. Additionally, there is a potential for these byproducts absorbed from Snus and ONPs to interact with the airways and lung microvasculature. Regarding other ways oral pouches/smokeless tobacco have been shown to impact the lungs, previous case studies have reported pulmonary aspirations of smokeless tobacco had induced multifocal airway obstructions and recurrent pulmonary infiltrations in the lungs of patients, the two reports cited suggest cases of ST-induced aspiration pneumonia [19, 24]. Smokeless tobacco -induced aspiration pneumonia and subsequent pulmonary inflammation can potentially be caused by direct contact between the airways and saliva that has come in contact with smokeless tobacco. Oral submucosal fibrosis is likely associated with pulmonary complications as seen with chewing tobacco. Studies have shown that smokeless tobacco usage significantly reduces antioxidant activity in saliva and significantly increases the level of toxic metals in saliva, including heavy metals known to induce the formation of Reactive Oxygen Species (ROS) in lung epithelial cells [25–26]. Additionally, smokeless tobacco-induced pulmonary inflammation may also be caused by regurgitated gastric stomach acid coming into contact with the airways; this occurs due to the nicotine absorbed into the bloodstream from oral pouches/ smokeless tobacco increasing the possibility of lower esophageal sphincter relaxation [27, 28].

Likewise, due to the potential of smokeless nicotine-pouch products to negatively impact lung function, the limited number of relevant studies, and the increasing popularity of ONPs in the U.S., we have conducted a study involving the analysis of changes in inflammatory cytokine release, ROS production, and cytotoxicity in bronchial/lung epithelial cells exposed to smoke-free nicotine-based pouch extract. Unlike previous studies conducted, which include analyses of cytotoxicity and cellular ROS among bronchial epithelial cells exposed to a variety of flavored ONPs, our study includes analyses using four of the most widely sold brands of ONPs containing TFN in the U.S., as well as analyses of inflammatory cytokines levels among bronchial epithelial cells exposed to these ONPs. Our study is the first to attempt to classify/categorize and elucidate how the usage of ONPs may impact lung epithelium. We employed the human bronchial epithelial cells BEAS-2B and 16-HBE cell lines to analyze cytotoxicity, cellular ROS, and cytokine release after exposure to ONPs. More specifically, our study utilizes LDH assays for examining cellular cytotoxicity, CellROX green assay for measuring ROS, and enzyme-linked immunoassays (ELISAs) for measuring levels of inflammatory cytokines, IL-6 and IL-8.

## MATERIALS AND METHODS

### Classification/Categorization of Oral Nicotine Pouches (ONPs)/products

The pouches were procured from local vendors based in Rochester, NY, USA, and are sold publicly with age restrictions. The classification/categorization of the extracts of different flavored ONPs was carried out based on flavor categorizations and nicotine concentrations (**Table 1 and Figure 1**). Various nicotine strengths, i.e., ranging from 3 mg to ~8 mg/pouch, are found in these pouches; these pouches also contain various levels of moisture content and alkalinity. ONPs generally contain sweeteners, flavorings, food grain fillers, and plant-based fibers (cellulose). The flavors and flavor categories of ONPs belonging to the most widely sold brands in the U.S are given in the wheel diagram (**Figure 1**). The flavors of ONPs made by **Rogue** (Swisher) include Wintergreen, Peppermint, Spearmint, Berry, Apple, Honey Lemon, Mango and Cinnamon. ONP flavors from **ON!** include Wintergreen, Cinnamon, Citrus, Coffee, Berry, and Original. ONP flavors from **Velo** formerly REVEL brand include Berry, Cherry, Cinnamon, Citrus, Coffee, Dragonfruit, Mint, Wintergreen, Peppermint, Cream, Vanilla, and Spearmint. ONP flavors from **Zyn** (Swedish Match) include Coffee, Cinnamon, Wintergreen, Spearmint, Citrus, Peppermint, Cool Mint, Original/Smooth Tobacco (unflavored or tobacco flavored). The **NIIN** ONP flavors include Wintergreen, Spearmint, Cool mint, Citrus chill, and Cinnamon. The **FRE** ONP flavors include Sweet, Lush, Wintergreen, and Mint. The **Killa** ONP flavors include cold mint, spearmint, Dutch cold, Watermelon, Blueberry, and Apple. The **Nordic Spirit** ONPs include Mint, Spearmint, Wild berry, Mocha, and Elderflower; ONP flavors from **Zonex** include Cold blast (mint and peppermint), Berry, and Breeze. The flavors of **Lyft** ONPs include Ice cool, Mint, Freeze X-Strong, Cool Air, Blueberry, Lime, Berry Twist, Blonde Roast, Melon, Strawberry, Licorice, and Tropic. ONP flavors from **Dryft** (Kretek) include Blackberry, Cinnamon, Citrus, Coffee, Dragon Fruit, Peppermint, Spearmint, and Wintergreen. Mint/mentholated flavors and fruit flavors of ONPS were analyzed in this study as these are two of the most widely sold ONP flavors in the U.S, according to one study using retail scanner data to assess nicotine pouch sales in the U.S from 2016-2020 [11].

**Table 1:** Flavor and nicotine concentration(s) of Smoke-free Nicotine Pouch-based products used.

| Product-type | Brand | Flavor | Nicotine-type | Nicotine Concentration (mg) |
| --- | --- | --- | --- | --- |
| ONP | ON | Original | TFN | 8 |
| ONP | Rogue | Mango | TFN | 6 |
| ONP | Velo | Black Cherry | TFN | 7 |
| ONP | Zyn | Cool Spearmint | TFN | 6 |
| Snus | SKOAL | Spearmint | TDN | 5 |

**Pouch product type, brand, flavor, nicotine-type (TDN or TFN), and nicotine concentration are listed in the table above.**

### Cells and culture conditions

The Human Bronchial Epithelial cell line (16-HBE) and BEAS-2B cell line (ATCC) were used in this study. 16-HBE cells were cultured in Dulbecco’s Modified Eagle’s Medium (DMEM) supplemented with 10% FBS and 1% antibiotic-antimycotic solution. BEAS-2B cells were grown in DMEM/F12 media with 10% FBS, 15μM HEPES, and 1% antibiotic-antimycotic solution. Cells were maintained at 37°C and 5% CO_2_ in a humidified atmosphere and used for the experiments. Passages below 10 were selected, and when the sufficient density was reached, cells were seeded at 250,000 cells per well in 48-well plates with 500 μL of complete DMEM media. Cells were incubated overnight in low serum-containing media (FBS 0%) to lessen undesired stimulation of the cells and the cytokine background levels. Serum starvation permitted us to compute subdued alterations in cytokine levels because of the treatment of interest.

### Extraction of Oral Nicotine Pouches

The extraction of Nicotine Pouches in PBS is shown in **Figure 2**. Extracts of oral nicotine pouches were created by incubating indicted pouches in PBS (1:10 w/v) for 1 h on a shaker (500 rpm) at 37°C. Extracts were centrifuged and then filtered through 0.45-micron sterile filters and denoted 100% for treatments [29–32]. The aliquots of 100 μL per extract were frozen at −20°C for experimental use.

**Figure 2:**
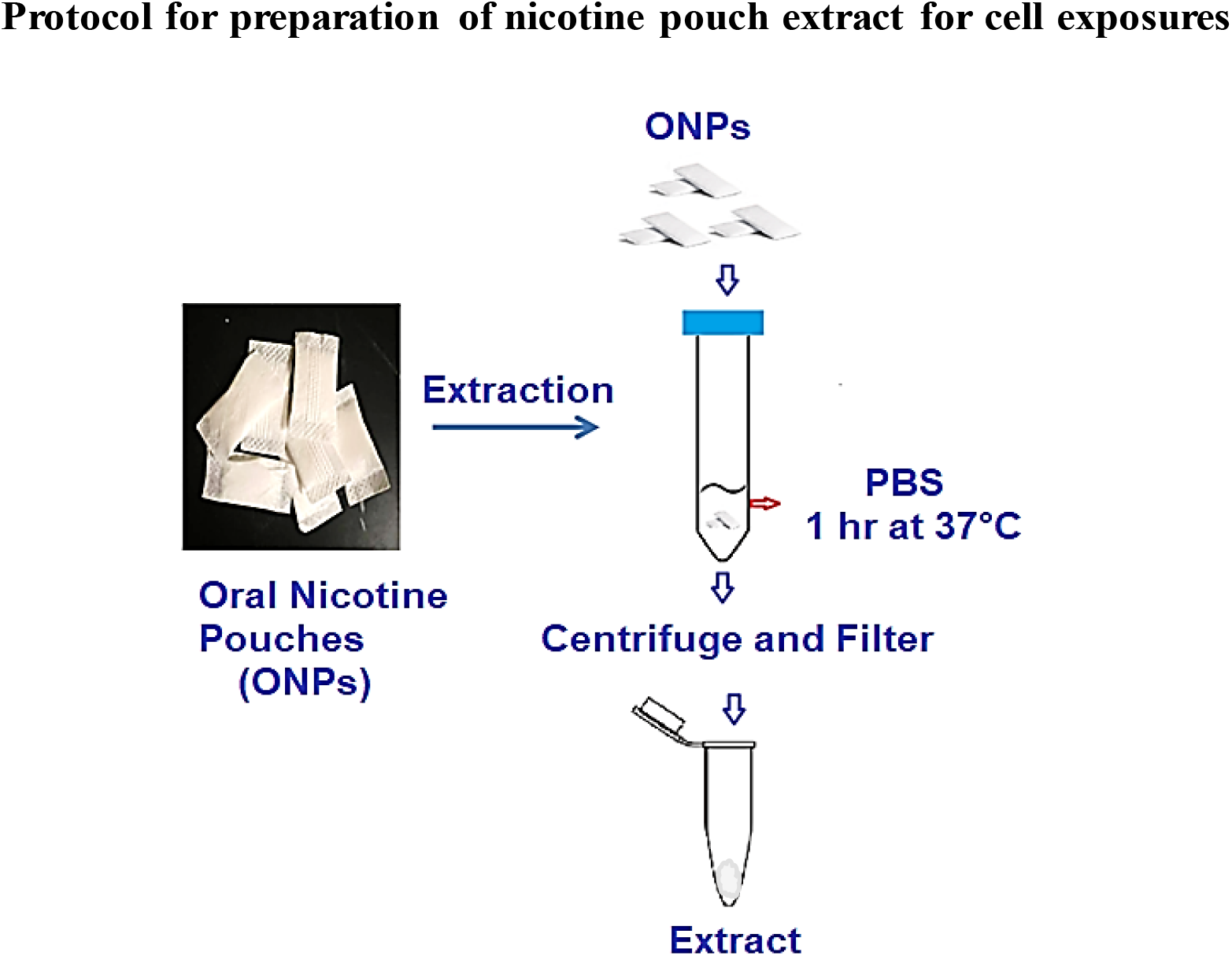
A Protocol for preparation of nicotine pouch extract for cell exposures. Extracts of oral nicotine pouches were prepared by incubating pouches in PBS (1:10 w/v) for 1h on a shaker (500 rpm) at 37°C. Extracts were centrifuged and then filtered through 0.45 micron sterile filters and estimated 100% for treatments. The aliquots of 100 μL per extract were frozen at −20°C for experimental use.

### Cell Treatments with ONP extracts and Conditioned Media Collections

Serum-deprived cells were treated with different flavored nicotine pouches. The BEAS-2B cells were treated with the Spearmint flavor, whereas the 16-HBE cells were treated with original (unflavored), and flavors including mango and black cherry. Respective treatments of nicotine pouches were used on the designated wells at varying concentrations (0.05%, 0.1%, 0.25%, and 1% in triplicates) [31,32]. To minimize cell death when assessing for cellular ROS and cytokine release, 24 hours post-treatment, the conditioned media was collected by centrifuging 16-HBE cell suspension at 1,000 rpm for 5 min and centrifuging BEAS-2B cell suspension at 1,000 rpm for 5 min. Subsequently, collected supernatants were frozen at −20°C to assess cytokine levels. The viability of the cells was measured by re-suspending the cells in DMEM using the acridine orange (AO) and propidium iodide (PI) staining for evaluating the live and dead cell concentration as a percentage (for seeding) via an automatic cellometer.

### Lactate Dehydrogenase (LDH) Cytotoxicity assay

Quantification of Lactate Dehydrogenase (LDH) release was used to assess the levels of cytotoxicity induced by exposure to extracts of ONPs and Snus. Following treatment of the nicotine pouches, the culture medium was aspirated and centrifuged at 1000 rpm for 5 min to obtain a cell-free supernatant. The activity of LDH in the medium was determined using a commercially available kit (Roche) per the manufacturer’s instructions. Aliquots of media and the required reagents were mixed in a 96-well plate, and absorbance was recorded at 490 nm using amicroplate spectrophotometer system. The outcome was presented as a percentage of control values.

### ROS assay by CellROX Green

16-HBE cells were serum-deprived and treated with respective oral nicotine pouch extracts. After 4 hrs of incubation, cells were stained with CellROX reagents (Green) and Hoechst 33342 was used as nuclear stain and viewed in Cytation cell imaging multimode reader (Agilent) immediately. A broad spectrum of ROS was determined in living cells using the fluorogenic indicator CellROX. Probes for this were purchased from Invitrogen (Carlsbad, CA) and applied according to the manufacturer’s instructions. Images were captured on a Cytation cell imaging multimode reader (Agilent). Similar image acquiring times and settings for intensity were used for all images obtained.

### Inflammatory Response (IL-6 and IL-8) Assay

After cell treatments, conditioned media were collected after 24 hrs of treatment of different concentrations of nicotine pouches. Pro-inflammatory cytokine (IL-8) and (IL-6) release was determined using the IL-6 and IL-8 ELISA kit (Invitrogen) according to the manufacturer’s instructions (ThermoFisher).

## STATISTICAL ANALYSIS

Statistical analyses of significance were performed by the Student’s T-test and one-way ANOVA (Tukeys/Dunnets multiple comparison tests) when comparing multiple groups using GraphPad Prism 7 (La Jolla, CA). Data are presented as means ± SEM. P < 0.05 is considered statistically significant

## RESULTS

### Differential cytotoxicity among bronchial epithelial cells exposed to different flavored oral smokeless nicotine products

BEAS-2B cells were exposed to different concentrations of extracts isolated from spearmint flavored Snus (SKOAL) and ONPs (Zyn), subsequently, the cytotoxicity of the flavored nicotine pouches was assessed through collecting culture media exposed to 24 hrs of treatment with a respective extract. The untreated group was considered the control group. Among the tested spearmint products, minimal LDH release was observed at 0.05% concentration, whereas exposure to the 0.1% and 0.25% concentrations exhibited significant levels of LDH release (p < 0.01) (**Figure 3A**). Specifically, both the % LDH release values of BEAS-2B cells exposed to 0.05% Skoal Spearmint extract and the 0.05% Zyn Spearmint extract did not significantly differ from the corresponding control (p>0.05). Additionally, both the % LDH release values of BEAS-2B cells exposed to 0.1% (v/v) Skoal Spearmint extract and 0.1 % (v/v) Zyn Spearmint extract did significantly differ from the % of LDH released from the corresponding control (***p<0.001); similar results were seen amongst the 0.25% (v/v) spearmint-flavored oral nicotine pouch extracts. We further carried out other assays using 0.1% and 0.25% concentrations in light of these observations. Furthermore, 16-HBE cells were individually exposed to 0.25% and 1% extracts of ON! original, Rogue Mango, or Velo Black Cherry ONPs. Differential LDH release was seen among all the different flavors of ONPs exposed to 16-HBE cells (**Figure 3B**).

**Figure 3A-B:**
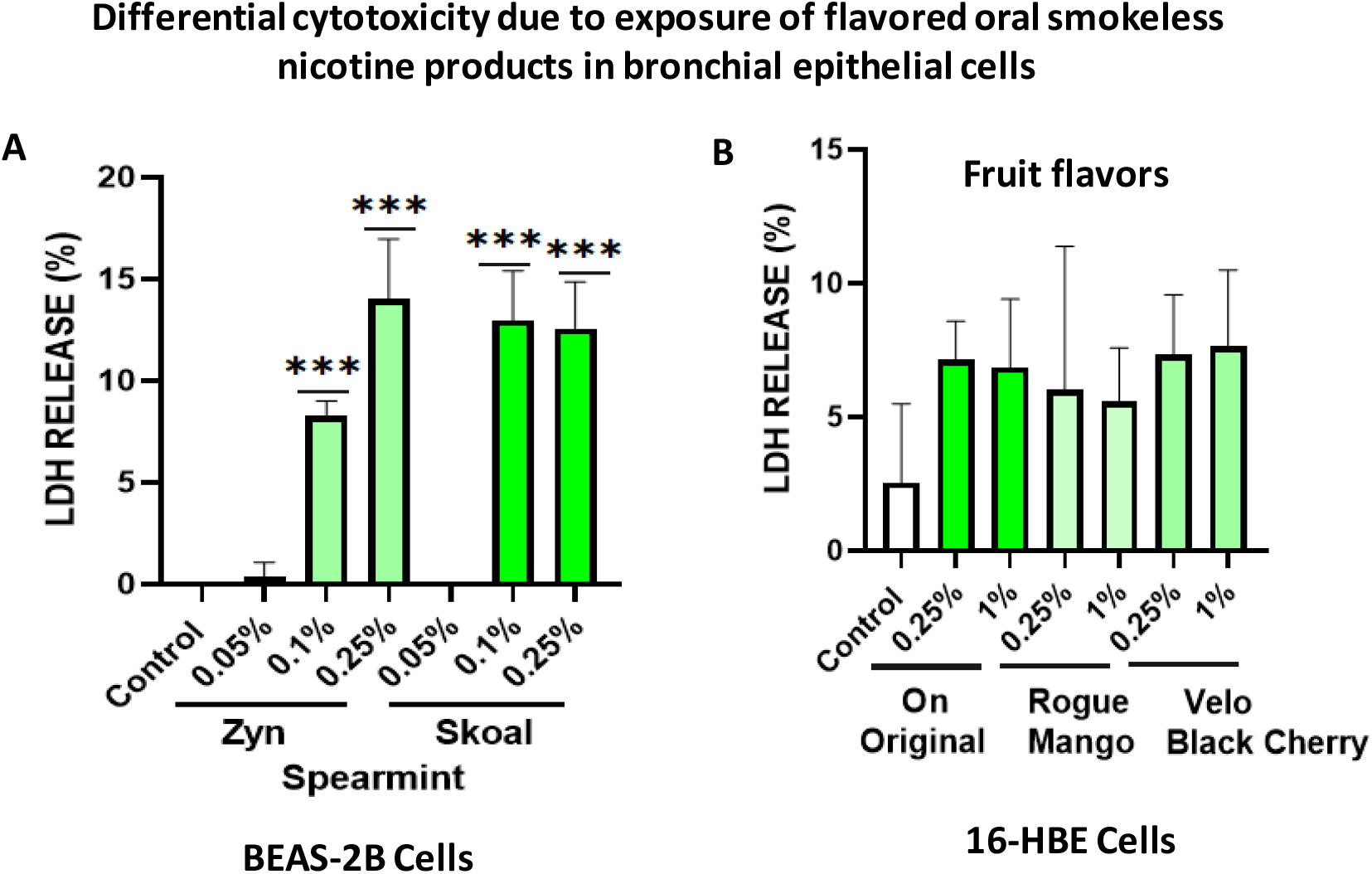
Differential LDH release (Cytotoxicity) by Oral Smokeless Nicotine Products. **A:** BEAS-2B cells were treated with different doses of spearmint flavored oral product extracts from two different brands (Zyn: ONP, Skoal: Snus). Control: Untreated cells. Following 24h exposure, conditioned media was used for Lactate Dehydrogenase (LDH) assay. Data represented as Mean ± SEM, ***p<0.001 compared to control. n = 3. **B:** 16-HBE cells were treated with different flavors of the oral product extracts of different brands. Following 24h exposure, conditioned media was used for LDH assay, n=3.

### ROS production in human bronchial epithelial cells

The level of ROS production related to the fluorogenic probe was assessed in 16-HBE cells treated with the flavored ONPs of interest. Cells were stained with CellROX reagents (Green), counterstained with Hoechst 33342 and viewed in Cytation cell imaging multimode reader (Agilent) immediately. CellROX green reagent is a fluorogenic probe that measures oxidative stress in live cells; these cells exhibit bright fluorescence upon ROS oxidation. As shown in **Figure 4**, 16-HBE cells treated with extracts of the original, mango, and black cherry-flavored ONPs showed higher levels of ROS production compared to the control-treatment, albeit these differences were not significant.

**Figure 4:**
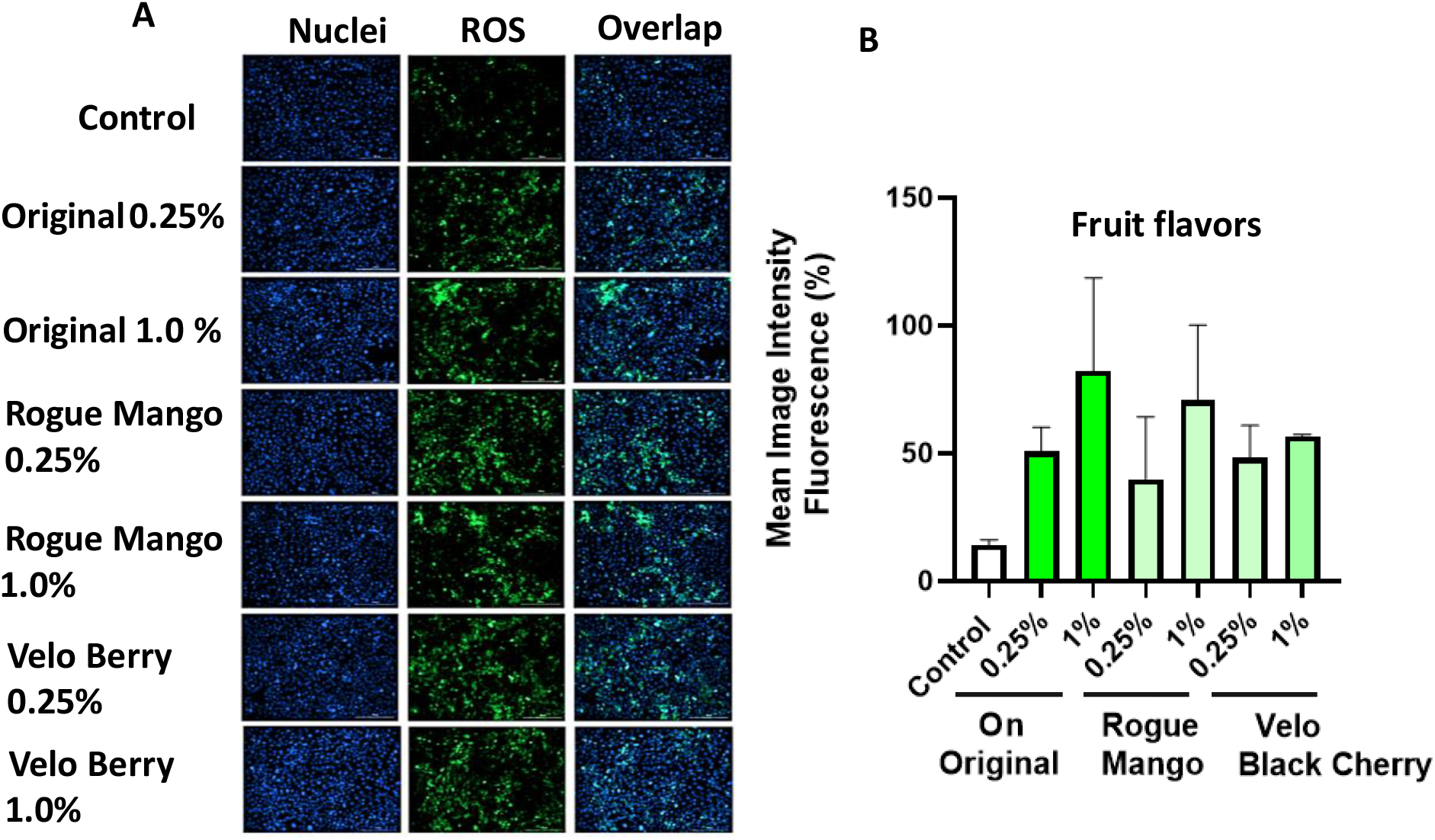
ROS release from 16-HBE cells after treatments with different flavors of oral nicotine pouches. 16-HBE cells were treated with different doses of ON Original, Rogue Mango or Velo Black Cherry flavored nicotine pouch extract and incubated for 4h. Following 4h exposure, cells were exposed to a CellROX assay. Images show CellROX (green) counterstained with Hoecht 33342.**A.** Fluorescence images display the release of ROS and cell nuclear morphology (blue) in 16-HBE cells treated with extracts isolated from different flavored ONPs at two different concentrations (0.25% and 1.0%). **B.** Quantitative analysis of fluorescence. Data represented as Mean ± SEM, Control (untreated cells), n = 3.

### Inflammatory mediator response due to flavoring nicotine oral products in bronchial epithelial cells

Through treating BEAS-2B and 16-HBE cells with extracts of flavored ONPs and Snus, ONP-induced inflammatory cytokine responses in bronchial epithelial cells were assessed; specifically, this was done through measuring IL-8 and IL-6 concentrations in conditioned media. BEAS-2B cells were treated with spearmint flavored Snus and ONPs across two concentrations, 0.1% and 0.25%. Treatment with the spearmint ONP (Zyn) led to significant increases in IL-8 levels in both dose conditions (p<0.05) (**Figure 5A**). Similar results were found for IL-6 release when the cells were treated with different concentrations of Zyn (unpublished observations). When analyzing IL-6 patterns among 16-HBE cells exposed to extracts of the ON, Rogue, and Velo ONPs used, across both concentrations (0.25% and 1.0%), we see that IL-6 levels were not significantly different from those of the control (**Figure 5B**).

**Figure 5:**
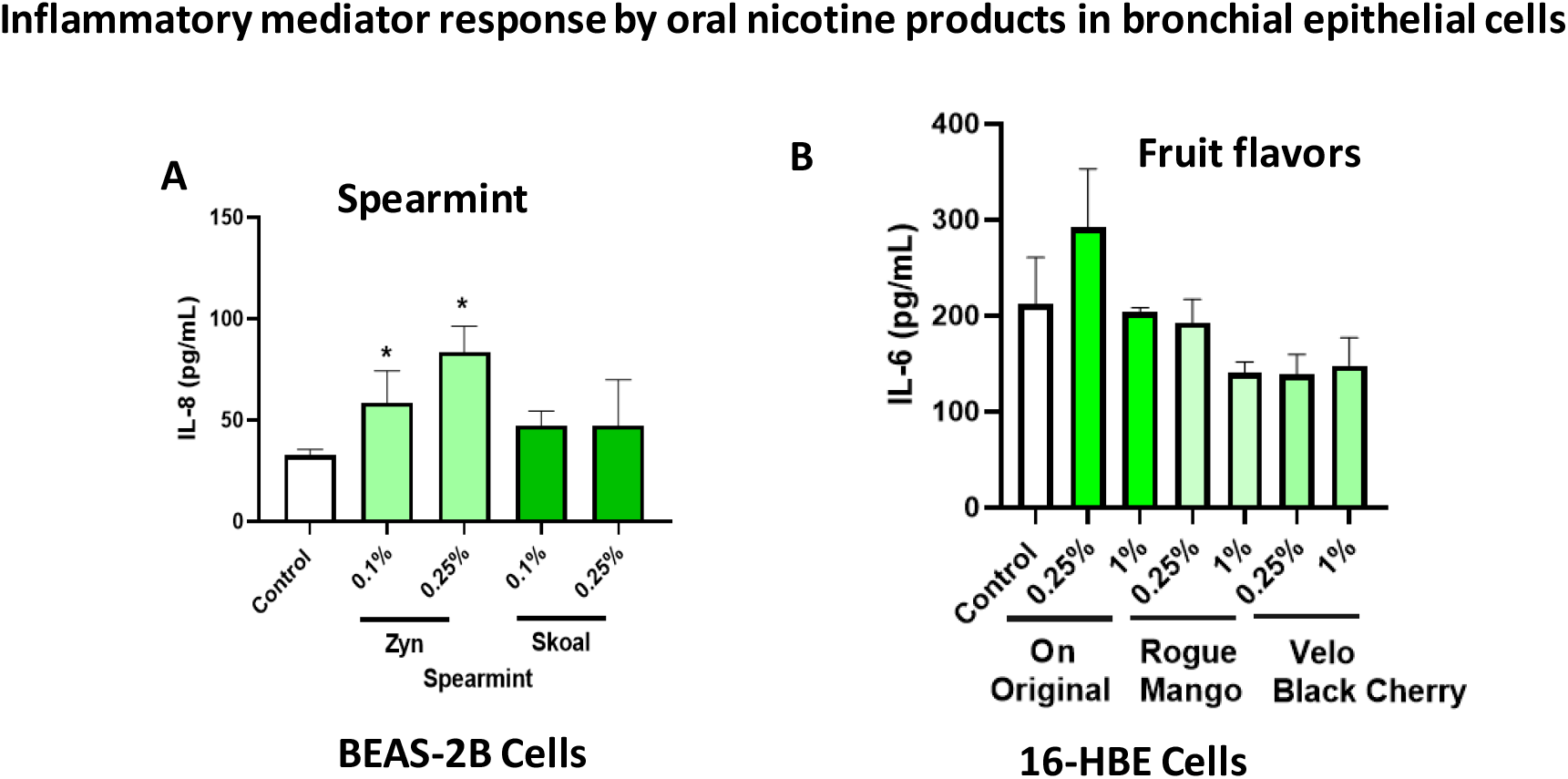
IL-8 and IL-6 release in response to flavored oral nicotine pouch extracts by bronchial epithelial cells. **A**: BEAS-2B cells were treated with the different doses of spearmint flavored nicotine pouch extracts from two different brands. Following 24h exposure, conditioned media was used for IL-8 assay. Data represented as Mean ± SEM, *p<0.05 compared to control (untreated), n = 3. **B:** 16-HBE cells treated with different oral nicotine pouch flavors; ON original, Rogue Mango or Velo Black Cherry and incubated for 24 hrs. Following 24 h exposure, conditioned media was used for IL-6 assay.

## DISCUSSION

With the prevalence of both ONP and Snus usage and the emerging popularity of flavored ONPs in the U.S., there is a crucial need better to understand the oral and pulmonary health effects of smoke-free nicotine-pouch-based products. Additionally, to discern the efficacy of oral-nicotine pouches as a potential nicotine replacement therapy product (NRTP) for those trying to quit vaping, studies that involve analyzing biomarkers of chronic e-cigarette use and e-cigarette- or vaping-associated lung injury (EVALI) using the most popular-brands of ONPs will be crucial. Such biomarkers include inflammation, oxidative stress (oxidant generation), mitochondrial dysfunction, and cell death among airway/bronchial epithelial cells. Our study sought to better elucidate how using ONPs may impact respiratory cellular effects. To do this, we assessed the levels of proinflammatory cytokine release, ROS production, and cytotoxicity in BEAS-2B exposed to extracts of ONPs of various volume concentrations (v/v%). Our study used four of the most widely sold flavored oral nicotine pouch brands in the U.S. (Velo, Rogue, ON!, and Zyn). Additionally, our study involved comparative analyses of inflammation, ROS production, and cytotoxicity among BEAS-2B cells between identically flavored ONPs across multiple extract volume concentrations.

Our data suggest that differences in the level of extract-induced cytotoxicity on human bronchial epithelial cells between flavored ONPs are dependent on volume concentrations (v/v%) of the extract used with the artificial saliva produced. Regarding the results of our cytotoxicity assay, specifically the % LDH release levels in cells exposed to extracts from the spearmint-flavored pouches used in the study, we found that differences in cytotoxicity between identical-flavored Snus and ONPs vary depending on the volume concentration of extract used (0.05, 0.1, and 0.25%). Part of our cytotoxicity data, specifically the % LDH released from BEAS-2B cells treated with Skoal Spearmint and Zyn Spearmint extracts, align with the finding of another study that utilized cytotoxicity assays using human lung epithelial cells exposed to Snus and oral nicotine-pouches by Bishop *et al*. [29]. Using human lung epithelial cells (H292) exposed to oral nicotine pouch extracts of various volume concentrations, Bishop *et al*. had shown that cells exposed to extracts from ONPs had shown less cytotoxicity than those exposed to extracts of Snus [30].

Regarding other comparisons between our cytotoxicity data and that compiled within other similar studies, the results of the LDH assays conducted in this study differed from that of the findings of a recent similar survey, East et al. 2021, which had similarly utilized bronchial epithelial cells exposed to the extracts of Snus and ONPs. East et al. 2021’s findings suggest that regardless of flavor or extract volume concentration, TDN-containing ONPs are less cytotoxic than Snus [30]. Specifically, East et al.’s cytotoxicity assays suggest that LYFT-Revel brand ONPs (now sold under the Velo brand) are significantly less cytotoxic across multiple nicotine concentrations and flavor types than Snus [30]. In our study, the results of the cytotoxicity assays conducted suggest that the difference between spearmint-flavored Skoal and Zyn extracts in inducing cytotoxic effects in BEAS2-B cells is negligible. However, the difference between our and East et al. 2021’s findings may be attributable to the differences in how our cytotoxicity assays were conducted. Unlike the comparative analyses in cytotoxicity between Snus and ONPs conducted by East et al. 2021, those within our study included Snus and ONPs of a single flavor variety (spearmint). Additionally, the single type of snus product analyzed in East et al.2021 was the CORESTA Smokeless Tobacco Reference Product (CRP1.1), an unflavored/tobacco flavored Swedish-style snus product [30]. Studies have shown that users of ONPs are exposed to lower levels of toxic compounds than users of Snus [4]. Additionally, unlike East et al. 2021, which only used TDN-containing ONPs, our study only used TFN-containing ONPs [30]. We further showed dramatic cytotoxicity in oral epithelial cells by these flavored ONPs (unpublished observations), this work is ongoing in our laboratory. To corroborate the findings of cytotoxicity, it would be interesting to determine the osmolarity and compare with pH values, and nicotine content (free or protonated forms) of the pouches/extracts or in saliva.

Regarding the release of ROS in response to flavored ONPs from human bronchial epithelial cells, our findings showed a differential response with higher levels of CellROX staining in all the treatment groups. 16-HBE cells treated with extracts of the original (unflavored tobacco), mango, and black cherry-flavored showed a differential and slightly higher levels of ROS production when treated with higher doses of extracts by fruity ONPs. Ongoing work with different flavored ONPs along with mint vs fruity (**Figure 1**) will differentiate the cellular ROS responses including the toxicity on redox homeostasis and mitochondrial function.

Regarding the results of proinflammatory mediators, our findings differed from one study that investigated pro-inflammatory cytokine responses due to exposure to oral pouches/smokeless tobacco [33]. To further explain, Zutsi *et al*., via quantitative analyses of serum inflammatory cytokine markers like tumor necrosis factor-alpha (TNFα), interleukin IL-6, and interleukin IL-1β, found that among healthy adults, chronic use of oral products is associated with subclinical systemic inflammation [33]. However, in our analysis of pro-inflammatory cytokine levels, we found that regardless of extract volume concentration (0.1, and 0.25%), BEAS-2B cells exposed to Snus (Skoal Spearmint) did not produce significant levels of IL-8. However, this difference between the findings of our study and that of Zutsi *et al*. may be due to the differences in how our inflammatory cytokine analyses were conducted; our study measured the level of a proinflammatory cytokine among cultured BEAS-2B cells while Zutsi *et al*. had assessed proinflammatory cytokine levels using peripheral venous blood [33]. The reasons for this pro-inflammatory response by spearmint may be the presence of flavoring chemicals (e.g. flavor acetals and/or the presence of cooling agents in mint pouches (may be WS compounds), which requires further chemical analyses.

Overall, our data showed increased cytotoxicity, differential release of ROS, cytotoxicity, and cytokine (IL-6 and IL-8) release at the lowest concentration treatments at 4-24 hours with the most striking response by spearmint ONP, whereas ONPs containing original tobacco, mango, and black cherry showed higher levels of ROS release in these cells. These data suggest that flavored ONPs are not safe and likely to cause systemic and local toxicological responses during their chronic usage. Further studies are in progress to determine the oral and pulmonary toxicity of a variety of ONPs flavors and flavorants using *in vitro, ex-vivo, and in vivo* systems, including human periodontal health.

Regarding limitations in our study, our study did not involve the exposure of ONPs extracts using oral mucosal epithelial cells; instead, our study only used 16-HBE and BEAS-2B cells. Oral mucosal epithelial cells are the first mucosal epithelial cells that come in contact with oral nicotine products upon intraoral placement. Likewise, analyzing the levels of cytotoxicity, inflammation, and generated cellular ROS amongst oral mucosal epithelial cells exposed to oral nicotine-product extracts of various flavors and volume concentrations will be important in better understanding the harmful health effects of oral nicotine pouch usage [34]. Studies have shown that regular use of pouches/smokeless tobacco products induces cytological changes in the oral mucosa [34, 35], e.g., oral submucosal fibrosis is associated with pulmonary complications in smokeless product users [36]. Additionally, studies have shown regular pouches/smokeless tobacco usage can induce oxidative stress, inflammation, and apoptosis in oral keratinocytes [37–39]. Similar studies conducted involving ONPs have so far only used two types of cells in the oral cavity, human oral fibroblasts (HGF) and human gingival fibroblasts (HGF) [29,30]. However, no studies utilizing the exposure of oral epithelial cells or 3D EpiOral epithelium to nicotine pouch extracts for analyses of proinflammatory cytokine response, cellular ROS, or cytotoxicity have been conducted.

Consequently, future studies involving analyses of cell responses to extracts from ONPs must include oral epithelial cells [40]. Similarly, a reference smokeless product (CRP1.1) and different nicotine strengths of ONPs (e.g. 3 mg/6 mg) may be used for comparing the results with each other. Despite the lack of data on cellular responses amongst oral epithelial cells, our findings suggest that there are variations in induced cytotoxicity, generated cellular ROS, and pro-inflammatory cytokine release in bronchial epithelial cells exposed to ONP extracts of various flavors and nicotine strengths. Additionally, these preliminary findings indicate the need for further evaluation of ONPs’ role in inducing systemic including oral and pulmonary health risks. Further studies are required to study the role of flavored ONPs with different vendors based on the same flavorings at different nicotine strengths which we recently identified based on perceptions [7]. The aforementioned experiment can minimize the role vendor/company plays as confounding factor in experiments focused on understanding differential flavored ONP-induced oral and pulmonary toxicity based on toxicity assessments among different flavored ONPs for tobacco regulatory science. At the same time, clinical studies are required to assess the toxicity of these emerging flavored ONPs on oral and pulmonary or systemic responses as shown previously using ENDS flavored products to better understand how systemic changes in inflammation affect the lungs and the pulmonary microvasculature [41–46].

## Acknowledgments

We thank Dr. Zidian Xie for providing assistance in preparation of flavor wheels. We thank Dr. Thivanka Muthumulage for his useful scientific and technical expertise.

## Funding

This research was supported by the TCORS Grant: CRoFT 1 U54 CA228110-01.

### Informed Consent Statement

Not applicable; no human subjects were involved.

## Author Contributions

Conceptualization, I.R.; methodology, I.R.; assay performance: SS, WC, JL, software, SS.; validation, WC and SS.; formal analysis, SS and WC.; investigation, SS and WC; re-sources, IR.; data curation, SS.; writing—original draft preparation, SS, S.Y, and I.R.; writing—review and editing, SS, S.Y., and I.R.; DL, preparation of schematics and conceptual diagrams; visualization, IR.; supervision, I.R.; project administration, I.R.; funding acquisition, I.R. All authors have read and agreed to the published version of the manuscript.

## Data Availability Statement

We declare that we have provided all the data in figures.

